# SELECTIVE INHIBITION OF SOLUBLE TNF ATTENUATES HIPPOCAMPAL NEUROINFLAMMATION AND PSD-95 EXPRESSION TO IMPROVE NEUROLOGICAL FUNCTIONS IN A RAT MODEL OF GULF WAR ILLNESS

**DOI:** 10.1101/2022.11.28.518204

**Authors:** Udaiyappan Janakiraman, Katelyn Larson, Nancy Nixon-Lee, Melissa Damon, Andrew Biscardi, Elisa Hawkins, Laxmikant S. Deshpande, Kirsty J. Dixon

**Affiliations:** Department of Surgery, School of Medicine, Virginia Commonwealth University, 1101 E. Marshall St, Richmond, VA, 23298, USA; Department of Neurology, School of Medicine, Virginia Commonwealth University, 1101 E. Marshall St, Richmond, VA, 23298, USA

**Keywords:** Gulf War Illness, diisopropyl fluorophosphate (DFP), soluble TNF, TNFR1, XPro1595, Neuroinflammation, synaptic plasticity, PSD-95, AMPA receptor, NMDA receptor, neurological functions, MRI, edema

## Abstract

**BACKGROUND:** Systemic inflammation is a major contributor to poor brain pathology across many disease conditions. Specifically, the upregulation of the pro-inflammatory cytokine TNF in the hippocampus activates its receptor TNFR1, reducing AMPA receptor trafficking to impair LTP and associated behavioral outcomes. Studies using animal models of GWI have shown both a chronic upregulation of TNF and impaired neurological function. Therefore, this study aimed to investigate whether selectively inhibiting only the soluble form of TNF (solTNF) that preferentially activates TNFR1 can reverse neuroinflammation to improve neuroplasticity and neurological function.

**METHODS:** GWI was induced in rats by treating with DFP (or vehicle) for 5 consecutive days. Six months later, the rats were treated with XPro1595 (or vehicle) for 2 weeks to selectively inhibit solTNF, after which they were subjected to a battery of behavioral tests (cognition, anxiety-related, depressive-like behavior, and neuropathic pain). MRI brain scans were performed, and the animals were euthanized for brain pathological analysis.

**RESULTS:** The hippocampus of the GWI rats had significantly increased neuroinflammatory levels, resulting in edema and reduced AMPA receptor trafficking to the post-synaptic membrane that collectively promoted impairments in memory, anxiety, depressive-like behavior, and neuropathic pain. However, treating the rats with XPro1595 in the chronic environment attenuated the neuroinflammatory response, that reduced edema and impaired AMPA receptor trafficking, allowing for improvements in all areas of neurological function.

**CONCLUSION:** Overall findings suggest that selectively inhibiting solTNF using XPro1595 reduces neuroinflammation, synaptic plasticity, and overall function when administered in the chronic setting of a rat model of GWI. This data supports the use of XPro1595 in Veterans with GWI.

## BACKGROUND

After the Persian Gulf War of 1990-1991, approximately 33% of U.S. Gulf War veterans returned with unexplained chronic physical and mental symptoms (1) (2) (3) (4) presently identified as Gulf War Illness (GWI). The exact pathophysiology of GWI remains unknown although recent studies suggest a possible role in neuroinflammation, including dysregulated activation of microglia and astrocytes (5) (6). Studies in rodent models of GWI using exposures to several chemical combinations have demonstrated neuroinflammation (7) (8) (9) (10) (11) (12) (13). GWI models utilizing DFP exposure have resulted in a distinct brain-wide neuroinflammatory response in the absence of evidence of brain damage (12), highlighting the capacity for these exposures to promote an underlying neuroinflammatory condition in GWI.

One of the key mediators of inflammation in many systemic chronic inflammatory and degenerative conditions is tumor necrosis factor (TNF) (14), (15), (16). TNF is a unique cytokine in that it is first produced as a transmembrane protein (tmTNF) that preferentially binds TNF receptor 2 (TNFR2: CD120b or p75/p80), but once cleaved from the cell membrane TNF exists in a soluble form (solTNF) that preferentially binds TNF receptor 1 (TNFR1: CD120a or p55/p60) (17). Even though both TNFR1 and TNFR2 can trigger some common signaling pathways (18), TNFR2 activation generally promotes beneficial outcomes such as cell survival, induction of neurogenesis, and promotion of CNS autoimmunity (19), (20), (21), while TNFR1 activity generally promotes harmful outcomes such as cell death, aberrant neuronal plasticity, and exacerbation of the existing inflammatory response (19), (22), (23). Increased levels of several proinflammatory cytokines and inflammatory factors such as TNF have been reported in the rat brain following DFP exposure primed with CORT (24) and those expressing mood and cognitive impairment following exposure to GW neurotoxicants (5), (6).

In Veterans with GWI, neuroimaging and behavior profiling has been warranted to investigate these aforementioned changes in inflammatory responses and their relationship to GWI symptoms. However, there are no valid biomarkers for diagnosing and monitoring GWI pathology. Magnetic resonance imaging (MRI) is an increasingly popular imaging technique for investigating the brain’s neuronal lesions. Diffusion tensor imaging has been used to generate fractional anisotropy and apparent diffusion coefficient maps to identify symptoms such as restricted diffusion or edema (25), (26). Further, GWI-related chemical exposure and subsequent inflammatory response can potentially lead to chronic damage to the hippocampus, even compared to other brain structures associated with memory function (27).

These inflammatory related neuronal lesions detected on MRI could be impacting the brain’s neuroplasticity response. TNF is a known regulator of synaptic plasticity, both under physiological (29) and pathological (30), (31), (32) conditions. Synaptic plasticity has been studied in detail for hippocampal synapses, where changes in spine morphology and associated with changes in synaptic strength that are governed by the activation of N-methyl-D-aspartate receptors (NMDARs) and involve the trafficking of AMPARs between synaptic and extra-synaptic regions of dendrites (33) (34). Postsynaptic AMPARs mediate most of the fast-excitatory synaptic transmission in the central nervous system, which is thought to be one of the key cellular underlying cognitive functions, such as learning and memory (35) (36). An increased number of surface AMPARs leads to long-term potentiation (LTP) that promotes learning and memory, whereas the removal of surface AMPARs results in long-term depression (LTD) (35) (36). While there are little-to-no electrophysiological studies describing alterations in hippocampal synaptic plasticity or transmission in GWI models, data suggests that glutamatergic, GABAergic, and dopaminergic synaptic plasticity and transmission are impaired in GWI (37), (38).

In this study, we are using the novel ‘second generation’ biologic, XPro1595 as a tool for reducing neuroinflammation, preventing AMPAR trafficking, and improving neurological function in a rat model of GWI. XPro1595 selectively neutralizes solTNF that preferentially binds TNFR1. XPro1595 has been successfully used in many pre-clinical inflammatory disease models (39), (40), (41), (42), (43) with no known side effects, it can cross the blood brain barrier (BBB) (44) and has a half-life of 19.1 hours (40). Over the past 5 years XPro1595 has been used in a number of phase 1 clinical trials, with data suggesting that XPro1595 may be safe and well-tolerated (45), and able to reverse brain white matter inflammation and serum inflammatory markers (46), suggesting that it may be capable of reversing chronic inflammatory levels in Veterans with GWI and thus improve neurological functions.

## METHODS

### Study Summary

Male Sprague-Dawley rats (Envigo) approximately 300 g were used in this study. Half of the rats received DFP injections (called ‘GWI’ rats), while the other half received vehicle injections (called ‘Naïve’ rats) (Figure 1). After 6 months, half of each group were treated with XPro1595 (10 mg/kg, S.C. twice weekly) or vehicle (phosphate buffered saline) for 2 weeks. Groups were Naive+Veh, Naïve+XPro, GWI+Veh, and GWI+XPro. Following XPro1595 or vehicle treatment the rats were subjected to a battery of behavioral tests. Behavioral testing was performed using the least stressful to the most stressful tests, in the order indicated below. No two tests were carried out on the same day. At the completion of the behavioral tests, the brains were scanned using Magnetic Resonance Imaging, and the animals euthanized for brain pathological analysis.

**Figure 1:**
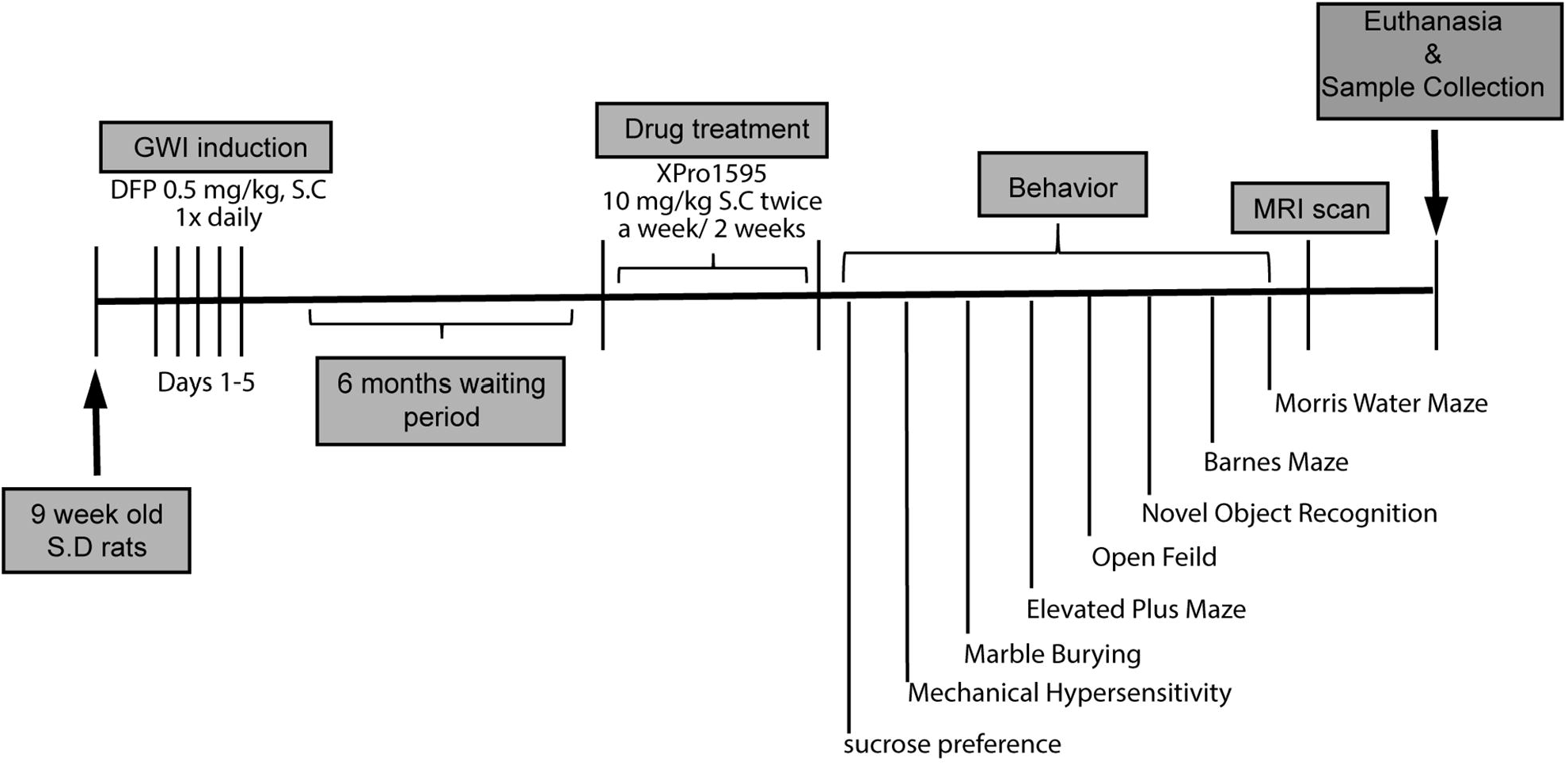
Chronological overview of the experimental design. Adult rats received DFP injections (or vehicle). After 6 months, half of each group were treated with XPro1595 (10 mg/kg, S.C. twice weekly, or vehicle) for 2 weeks, after which the rats were subjected to a battery of behavioral tests. At the completion of the behavioral tests, the brains were scanned using MRI, and the animals euthanized for brain pathological analysis. GWI = Gulf War Illness; DFP = diisopropylfluorophosphate; MRI = Magnetic resonance imaging.

### GWI Model

All animal use procedures were in strict accordance with the Virginia Commonwealth University Institutional Animal Care and Use Committee (protocol AM10039), and in accordance with NIH care and use of laboratory animals. Rats were housed two per cage, with a 12-hour light-dark cycle, and with ad libitum access to food and water, unless stated otherwise. Nine-week old rats were injected with DFP (0.5 mg/kg, S.C.; D0879, Sigma) or vehicle (ice-cold PBS) once daily for 5 consecutive days, according to (47) (48) (49) (50) (51) (52) (53).

### Immunohistochemistry

Rats were anaesthetized (4% isoflurane) and transcardially perfused with 300 mL of phosphate buffer saline (pH 7.2), followed by 300 mL of 4% paraformaldehyde. The brain was removed and incubated in 4% PFA for 2 hours, then cryoprotected in 15% sucrose for 24 hours, and then 30% sucrose for 1 week, prior to freezing. Sections (25 µm thick, coronal serial sections) were cut on a cryostat prior to storing at - 80 °C. Sections were permeabilized with 0.2% Triton X-100 in 2% fish gel for 2 hours at room temperature and immunohistochemically labelled with the primary antibody (1:200 rabbit anti-GFAP, Z0334 Dako; 1:200 rabbit anti-Iba1, 019-19741 Wako) overnight at 4 °C. Sections were washed 3 times in PBS, incubated in fluorescent secondary antibodies (1:500, Molecular Probes) for 90 min at room temperature, washed an additional 3 times in PBS, and coverslipped in mounting medium containing DAPI (ThermoFisher). The antibodies used in this study are listed in Supplementary Table S1.

### Golgi Staining

Rats were anaesthetized (4% isoflurane) and transcardial perfusion performed using approximately 30 ml of PBS, followed by approximately 100 ml of 4% paraformaldehyde. The brain was dissected and prepared according to FD Neurotechnology “Rapid Golgi stain” manufacturer’s instructions. 100 µm thick sagittal cryostat sections were cut according to the manufacturer’s instructions, and coverslipped in Permount mounting medium.

### Histological Analysis

Immunolabelled sections were photographed at 20x with equal exposure on a Zeiss AxioImager Z2 microscope, connected to a Zeiss Monochrome digital camera (AxioCam MRm) with Zeiss Zen Software (Blue Edition version 1.1.2.0). Glial reactivity within the ipsilateral hippocampus was determined by quantitating GFAP and Iba1 immunohistochemistry density, whereby images were converted to grayscale and thresholded using NIH ImageJ software (version 1.52a) to identify the area fraction of pixels positive for GFAP or Iba1-immunoreactivity. Values of each photograph per section were averaged, then averaged per animal, and then per group.

Golgi-stained sections containing the hippocampus were scanned for dendrites at least 200 µm in length in the CA1 region. These dendrites were photographed using a 100x oil objective on a Zeiss AxioImager Z2 microscope using Köhler illumination, connected to a Q Imaging color digital camera (Model 2000R-F-CLR-12, 12-bit) with Neurolucida software (version 2020.2.4). Photographs were quantitated to determine dendritic spine density, whereby the number of dendritic spines along a 200 µm long dendritic segment were counted and expressed as the number of spines per 200 µm.

### qPCR

Fresh hippocampal tissue was snap frozen in liquid nitrogen, and stored at -80 °C. From the frozen hippocampi, mRNA was extracted using the TRIzol method and converted to cDNA using a reverse transcription kit (Applied Biosystems), followed by mRNA expression being processed using SYBR green real-time PCR (Applied Biosystems). Inflammatory cytokine expression for TNF, IL-1β, and IL-6 was evaluated using Bio-Rad rat primers. Values were normalized to GAPDH expression and expressed as a fold change within the tissue, as compared to the control group (Naïve + vehicle).

### Whole Hippocampus Isolation

Rats were anaesthetized (4% isoflurane) and fresh hippocampi tissue was homogenized in RIPA buffer containing protease and phosphatase inhibitors and centrifuged at 20,000 x g for 20 min at 4 °C. The supernatant was collected and stored at -80 °C for Western blot analysis.

### Synaptoneurosome Isolation

Rats were anaesthetized (4% isoflurane) and fresh hippocampi were dissected (54). Briefly, the tissue was homogenized in a ‘homogenization buffer’ (Sucrose, 100 mM EGTA, 10 mM Tris, protease and phosphatase inhibitors (Roche), pH 8.1) at 4 °C. The homogenate was centrifuged at 2,000 x g for 5 min at 4 °C. The supernatant was centrifuged (MLA-55 fixed angle rotor) at 30,000 x g for 10 min at 4 °C. The pellet was resuspended in 400 μl of homogenization buffer, prior to the resuspended homogenate being layered on top of a 5-10.3% discontinuous Ficoll gradient and centrifuged at 45,000 x g for 20 min at 4 °C. The white layer in the 5-10.3% Ficoll interface that contains the synaptoneurosome was collected and centrifuged again with homogenization buffer at 30,000 x g for 10 min at 4°C. The pellet was collected, resuspended in 150 µl RIPA buffer containing protease and phosphatase inhibitors, and stored at -80 °C for Western blot analysis.

### Western Blotting

Protein concentrations were determined using the Bradford method (B6916, Sigma), samples were boiled for 5 min in 4x-Laemmli’s sample buffer (1610747, Bio-Rad), run on SDS-polyacrylamide gels (NP0321, Invitrogen), and transferred to PVDF membrane (1620177, Bio-Rad). Membranes were blocked in TBST solution (50 mM Tris-HCl, pH 7.5, 150 mM NaCl, and 0.1% Tween 20) containing 5% non-fat dry milk (Kroger supermarket) at 4 °C for 1 h, and then incubated overnight at 4 °C with anti-PSD-95 MAGUK scaffold protein (1:200; NeuroMab, clone K28/43), anti-phospho-AMPAR1 (S845) (1:1000; PPS008, R&D systems), anti-AMPAR1 (1:1000; AB1504, Millipore), anti-Synaptophysin (1:1000; MAB368, Chemicon), anti-EAAT2 (1:1000; sc-365634HRP, Santa-Cruz Biotechnology), anti-phospho-NMDAR1 (Ser896) (1:1000; PA537589, Invitrogen), anti-NMDAR1 (1:1000; MAB1586, Millipore) and β-actin (1:3000; sc-47778HRP, Santa-Cruz Biotechnology) in 3% BSA (BP1600, Fisher). After washing in Tris buffered saline containing 1% tween-20, membranes were incubated with secondary antibodies diluted in 3% BSA. HRP was visualized using chemiluminescence in a Bio-Rad ChemiDoc machine, and quantified band measurement using Bio-Rad Quantity One (version 4.6.6) optical density calculation (arbitrary units). Protein of interest expression levels were normalized to beta-actin loading control levels. For TNFR1 protein levels two different Western blots were run, with each blot having n = 2 per group (totaling n=4 per group). For synaptoneurosome protocol optimization one Western blot was run, with n = 2 samples per group (see supplementary section), while for the synaptoneurosome experimental conditions for each protein of interest, we ran either two or three different Western blots, with each blot having either n = 3 or 2 samples (respectively) per group, totaling n = 6 per group. On each membrane, each sample was normalized to beta actin levels, prior to averaging the n=6 per group across the membranes. Ratio values were obtained by dividing AMPA receptor levels by PSD-95 levels.

### Magnetic Resonance Imaging

MRI will be performed on a 7-Tesla 30-cm horizontal bore magnetic resonance imager (Bruker Advance III). The unit is equipped with a 12-cm inner diameter actively shielded gradient coil insert driven by high-output linear amplifiers allowing a maximum gradient strength of 600 mT/m. Radiofrequency (RF) transmission and reception are performed with a quadrature transmission RF coil of inner diameter 72 mm and an actively decoupled 4-channel parallel array RF coil specifically designed for mouse brain imaging. T1 relaxation maps will be acquired using a spin echo, echo-planar imaging sequence preceded by an inversion recovery preparation excitation. Individual images with inversion recovery times of 100, 180, 322, 578, 1038, 1863, 3343, and 6000 ms will be acquired. General acquisition parameters for the spin-echo images will include TR/TE of 7000/33 ms, matrix size of 128×128, 20×20 mm field of view, and slice thickness of 1.0 mm. T1 maps are then calculated using an exponential curve fitting algorithm. DTI scans will be acquired using a spin-echo echo-planar imaging protocol incorporating diffusion-sensitizing gradients. The diffusion-sensitizing gradients will be applied in 12 co-planar directions at 2 sensitizations, or b, values. The basic parameters for a DTI acquisition will be TR/TE of 3000/27 ms, matrix of 128×128, 20 × 20 mm field of view, 1.0 mm thick slices, and diffusion sensitizing b-values of 0 and 800 s/mm^2^. Quantitative assessments of data generated from T1, and DTI scans will be made by using the Image J software (Version 1.53t) to assess edema (DTI to calculate (∼ bregma -3.14 mm) edema, using T1 to demarcate area (∼ bregma -0.80 mm)). Region-of-interest was hand drawn on the calculated T1 Maps and DTI images. Pixel intensity was measured, and edema and lateral ventricles size were calculated.

### Behavioral Testing

#### Sucrose Preference Test (SPT)

Rats were habituated to two bottles (250ml sipper bottles; Allentown Inc.) in the cage lid for 5 days. The bottles were fitted with ball-bearing sipper tubes that prevented fluids from leaking. At the end of day 5 the rats were deprived of water but not food overnight. On the following day, two bottles were again introduced to the cage top for a period of 1 hour, with rats having the free choice of drinking either the 2 % sucrose solution in regular tap water, or 100% regular tap water. The volume of fluid and the weight of the bottle were measured before and after the test. The amount of sucrose intake was calculated as a percentage of total fluid intake.

#### Hindpaw Mechanical Hypersensitivity

Rats were placed on top of steel mesh (0.5 x 0.5 inch grid squares; 492398 Lowes), and allowed acclimatize for 4 minutes. Hindpaw hypersensitivity was assessed using mechanical von Frey filaments (Bioseb) starting with the lightest filament size (2.04 g) and sequentially testing up through the filaments until a positive test was established (hindleg rapid withdrawal or licking of the paw), which was recorded for each animal.

#### Marble Burying (MB)

Plastic cages (45 cm long × 23 cm wide × 20 cm high) were filled with 5 cm of fresh bedding, and black marbles (25 marbles; 2 cm diameter) with a slight metallic sheen were arranged on top of the bedding in a regularly spaced grid in a triangle shape. The rats were placed in the center of the cage and allowed to freely roam for 20 min, after which they were returned to their home cage. For each rat, the number of buried marbles was recorded. Marbles are considered buried if at least 2/3 of the marble was submerged under the bedding.

#### Elevated Plus Maze (EPM)

The maze consisted of four arms (50 cm long x 10 cm wide), connected by a central square, 10 cm × 10 cm. Two of the arms were left open without walls, while 2 arms were closed on the sides and end (31 cm high black plexiglass walls). The maze was elevated 55 cm above the floor level and was set in a dimly lit room. The rats were placed at the center of the open and closed arms, facing the open arm, opposite the experimenter and allowed to explore the maze for 5 minutes. The video-tracking system ANY-maze (version 6.3; Stoelting Co.) was used to determine the percent of open arm entries (number of open arm entries / number of total arm entries x 100) and the percent of time spent in the open arm (open arm time / total test duration x 100). Anxiety index was also calculated (1 – [([open arm time/test duration] + [open arm entries/total number of entries])/2]) (55) (56) (57) (58).

#### Open Field

The rats were placed into the peripheral zone of an open field chamber (120 cm circular arena, with 30 cm high plexiglass walls) and their behavior was observed for 5 min (59). The ANY-maze software divided the arena into central (50 cm diameter) and peripheral zones, with time spent in the central zones analyzed.

#### Novel Object Recognition (NOR)

The Open Field arena described above was also used for the NOR test. Two identical objects (ceramic garden frog figurines, approximately 10 cm high) were placed on opposite sides of the arena, 20 cm from the edge. The rats were placed in the center of the arena and allowed to explore the space for 5 minutes, prior to being returned to their home cage. After 1 hour, one of the ‘old’ objects were removed and replaced with a ‘novel’ object (ceramic garden tree figurine, approximately 10 cm high), the rats were again placed in the center of the arena and allowed to explore for 5 minutes, prior to being returned to their home cage. The video tracking system ANY-maze data was analyzed for time spent exploring the old object, time spent exploring the novel object, and the discrimination index (time spent with the novel object / total object (old and novel) duration x 100).

#### Barnes Maze (BM)

The Barnes maze is a white circular platform (120 cm diameter, Med Associates), elevated 90 cm above the ground, with 18 holes (10 cm each in diameter) spaced evenly around the edge of the platform. The maze is brightly lit from above, but one of the holes has a box located underneath for the rat to escape the bright light (escape box). Visual cues were placed on the surrounding walls (colored star, circle, square and triangle) approximately 21 cm in height, placed 50 cm away from the maze. The acquisition phase consisted of 4 days of training (2 trials/day), with each rat being placed in the center of the platform and given 2 minutes to explore the platform and find the escape box, prior to being returned to their home cage. After 96 hours (day 8), the test phase rats were again placed in the maze’s center and allowed explored the platform for 120 seconds. ANY-maze software analyzed the time spent near (within 10 cm from) the target hole, time spent in target quadrant.

#### Morris Water Maze (MWM)

The MWM tank is a 6-foot wide circular galvanized steel stock feed tank (WTR62; Countyline) filled with water 21-inches deep. Approximately 500 ml white paint (447516, Valspar) was added to make the water opaque. The pool was divided into 4 quadrants, arbitrarily nominated North, South, East and West, and the escape platform (plexiglass circle 7 inches in diameter, covered in 5 layers of cheese cloth) was hidden 2 cm below the surface of the water in the center of the North quadrant. Visual cues were placed on the surrounding black walls (brightly colored star, circle, square and triangle) approximately 50 cm in height, placed 1 meter away and 50 cm higher than the tank. The ‘acquisition trials’ were performed on 5 continuous days, consisting of 4 trails per day, starting from each of the quadrants. If the rats did not reach the platform in 120 seconds, they were gently directed to the platform where they remained on top for 20 seconds, prior to being removed and returned to their home cage. On day 8, the hidden platform was removed and the ‘probe trial’ was performed. The rats were placed in the South quadrant and allowed swim for 30 seconds, prior to being removed and returned to their home cage. During the acquisition phase the rats escape latency (time taken to reach the platform) was determine using ANY-maze software, while during the probe trial the time spent in the target quadrant and number of entries into the target quadrant were analyzed.

### Statistical Analysis

All data were assessed for normality. Data were analyzed with GraphPad Prism 9. Group differences were calculated using one-way or repeated-measures (BM and MWM) ANOVA followed by Tukey’s post hoc test as indicated in the figures. P values smaller than 0.05 were considered significant. Data in figures are expressed as mean ± standard error of the mean. Only statistically significant outlier data points were removed.

## RESULTS

### GWI Promotes Sustained Inflammatory Cytokine Expression, Without Altering Cytokine Receptor Levels

To begin investigating the neuroinflammatory sequelae in rat model of GWI, major inflammatory cytokine mRNA expression was analyzed by qPCR in rat hippocampal tissue. qPCR analysis revealed significantly increased TNF expression in GWI+Veh rats (Figure 2A), as compared to the Naïve+Veh group. Conversely, treatment of the GWI rats with XPro1595 reduced TNF expression, such that expression levels were no longer significantly higher than that of the Naïve+XP and Naive+Veh groups. The same effect was observed for IL-6 (Figure 2B), whereby expression was significantly increased by nearly 30-fold in the GWI+Veh group, as compared to Naïve+Veh group, but was no longer significantly increased in the GWI rats treated with XPro1595. No changes in IL-1β (Figure 2C) gene expression were noted in any groups.

**Figure 2:**
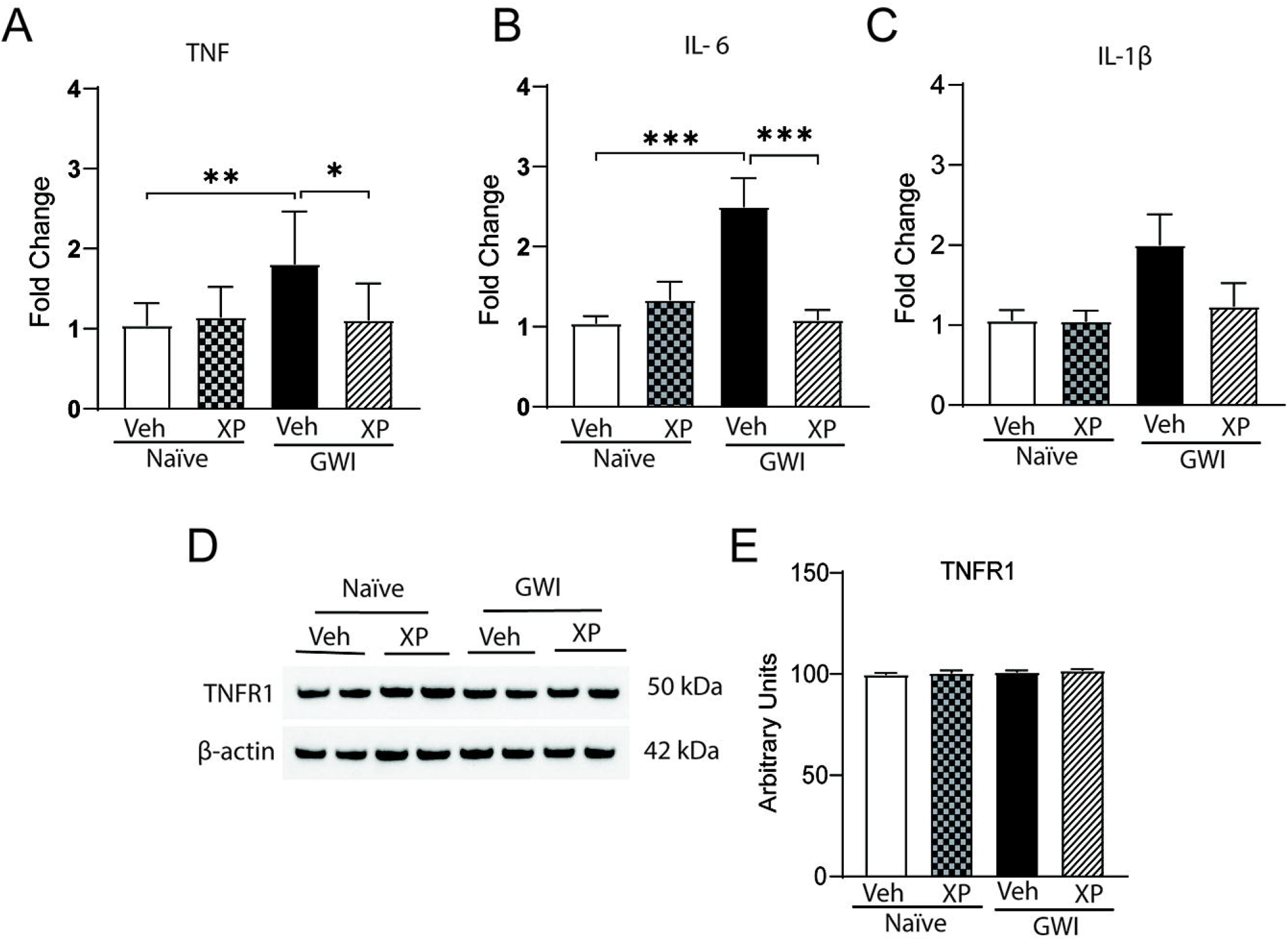
Six months after DFP exposure cytokine expression levels were quantitated. Hippocampal tissue dissected from rats in the GWI+Veh group showed significantly increased inflammatory cytokine gene expression (TNF and IL-6; **A&B**) compared to tissue from the control group (Naïve+Veh). There was a strong trend for the upregulation of IL-1β in the GWI+Veh group (**C**) that was not observed following XPro1595 treatment. Two weeks administration of XPro1595 reversed this effect by significantly reducing inflammatory gene expression back to baseline levels. Conversely, no changes were observed in hippocampal TNFR1 protein expression across groups (**D&E**). qPCR experiments used n=9 per group, while Western blotting used n=4 per group; * p = 0.05, ** p = 0.01, *** p = 0.001.

Western blotting was used to determine hippocampal TNFR1 expression levels. Analysis revealed that the vehicle-treated GWI group (GWI+Veh) did not have elevated TNFR1 expression compared to the vehicle-treated Naïve group (Naïve+Veh) chronically following DFP exposure (Figure 2D&E). Similarly, no changes were observed in the XPro1595-treated GWI group (GWI+XPro).

### GWI Promotes Astrocyte and Microglial/Macrophage Reactivity, which is Attenuated by Neutralizing solTNF

To determine whether an altered neuroinflammatory response impacted glial reactivity in our model of GWI, immunohistochemistry was used to semi-quantitate astrocyte (GFAP) and microglial/macrophage (Iba1) immunoreactivity levels and cell density.

Minimal GFAP immunoreactivity was observed in the hippocampal DG (Figure 3A&C) and CA1 (Figure 3B&E) regions in both of the Naïve rat groups. The GWI model (GWI+Veh) promoted sustained astrocyte reactivity, as observed by a significant increase in both GFAP immunoreactivity and number of GFAP-positive cells (Figure 3D&F) 6 months post-induction, as compared to the Naïve groups. Conversely, treatment with XPro1595 (GWI+XPro) attenuated this increase such that the level of GFAP immunoreactivity and GFAP-positive cell density was not significantly different to the Naïve groups.

**Figure 3:**
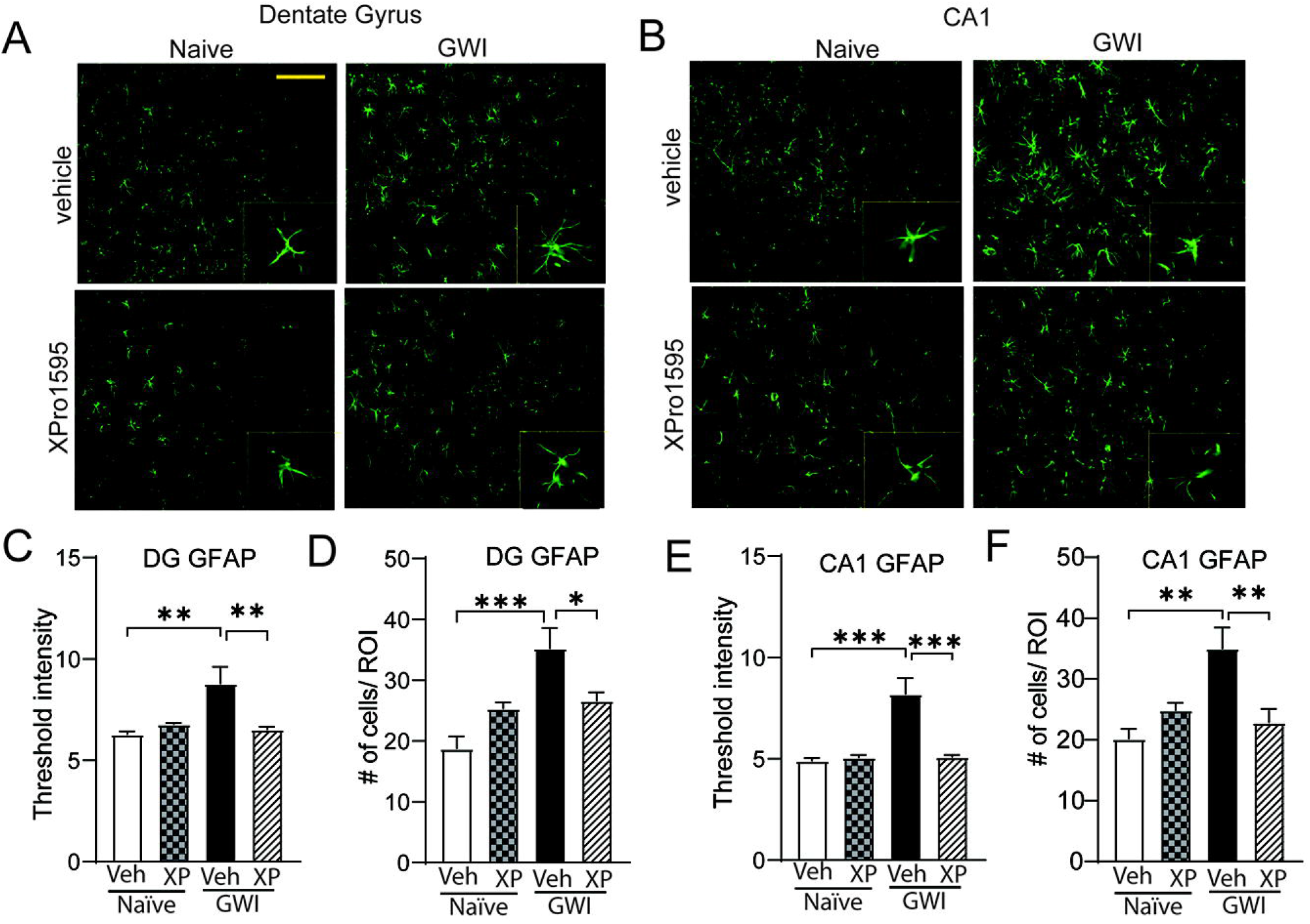
Astrocyte reactivity was assessed by quantitating hippocampal DG and CA1 GFAP expression (**A, B, C & E**) and the number of GFAP-positive cells (**A, B, D & F**) in the field of interest. In both hippocampal regions GFAP expression and number of GFAP-positive astrocytes was upregulated in the GWI+Veh group compared to the control group (Naïve+Veh), which was significantly reduced following XPro1595 treatment (GWI+XPro). n=5 animals per group; * p = 0.05, ** p = 0.01, *** p = 0.001; Scale bar in A = 200 μm.

Similar to astrocyte reactivity, minimal Iba1 immunoreactivity was observed in the hippocampal DG (Figure 4A&C) and CA1 (Figure 4B&E) regions in both of the Naïve rat groups. The GWI model promoted sustained microglial reactivity, as observed by a significant increase in both Iba1 immunoreactivity and number of Iba1-positive cells (Figure 4D&F) 6 months post-induction, as compared to the Naïve groups. Conversely, treatment with XPro1595 (GWI+XPro) attenuated this increase such that the level of Iba1 immunoreactivity and Iba1-positive cell density was not significantly different to the Naïve groups.

**Figure 4:**
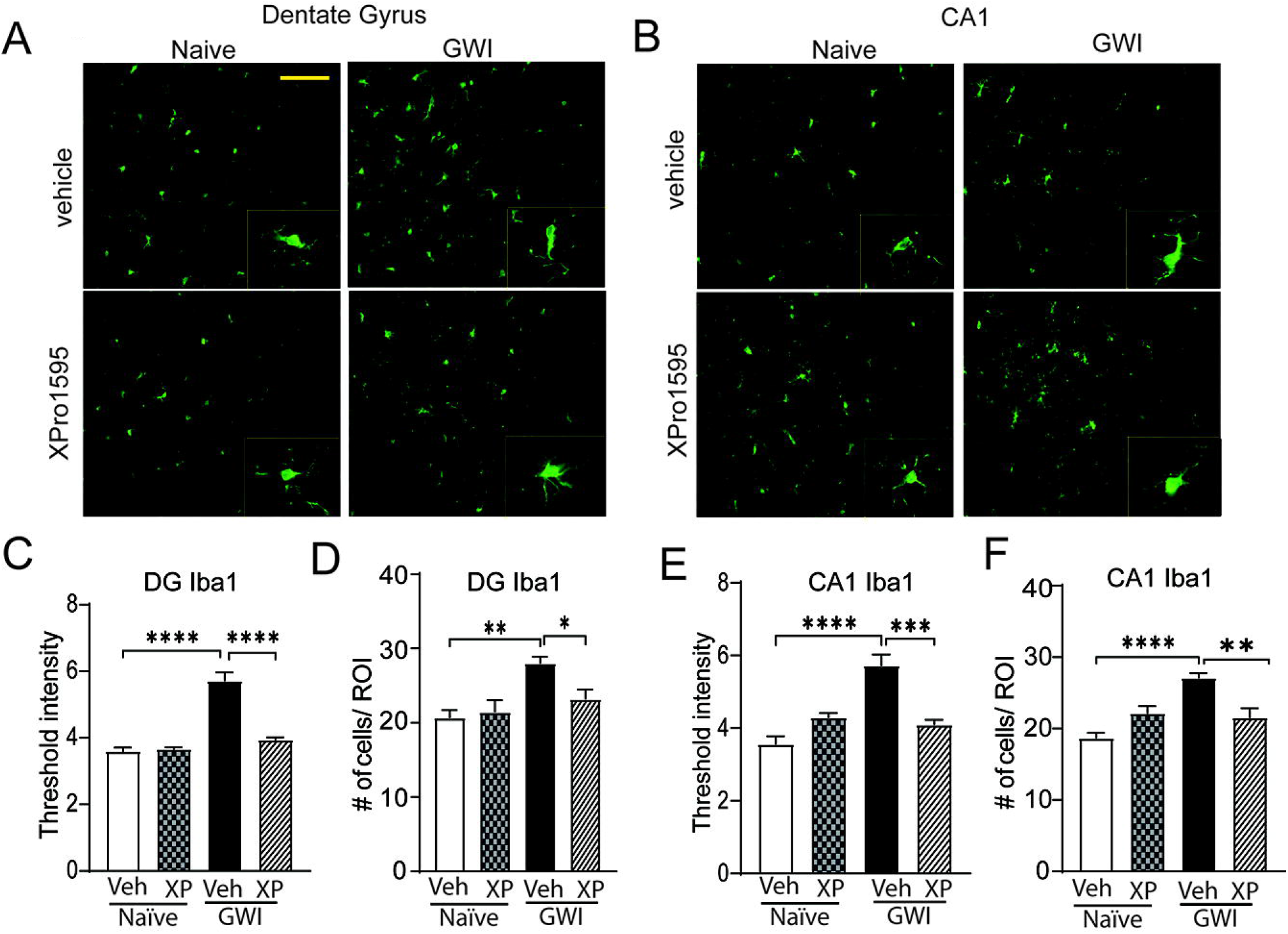
Microglial activation was assessed by quantitating hippocampal DG and CA1 Iba1 expression (**A, B, C & E**) and the number of Iba1-positive cells (**A, B, D & F**) in the field of interest. In both hippocampal regions Iba1 expression and number of Iba1-positive microglia was upregulated in the GWI+Veh group compared to the control group (Naïve+Veh), which was significantly reduced following XPro1595 treatment (GWI+XPro). n=5 animals per group; * p = 0.05, ** p = 0.01, *** p = 0.001, **** p = 0.0001; Scale bar in A = 200 μm.

### GWI Promotes Hippocampal Edema, which is Attenuated by Neutralizing solTNF

Given that inflammation can promote edema, the rat brains were scanned using T1 and DTI MRI (Figure 5A) to determine whether the increased the neuroinflammatory response increases hippocampal edema. Analysis of the DTI scans showed that edema is significantly increased in the hippocampus of GWI+Veh rats, as compared to the Naïve rat group (Naïve+Veh) (Figure 5B). In contrast, hippocampal edema levels in GWI rats treated with XPro1595 (GWI+XPro) were not significantly different to Naïve levels.

**Figure 5:**
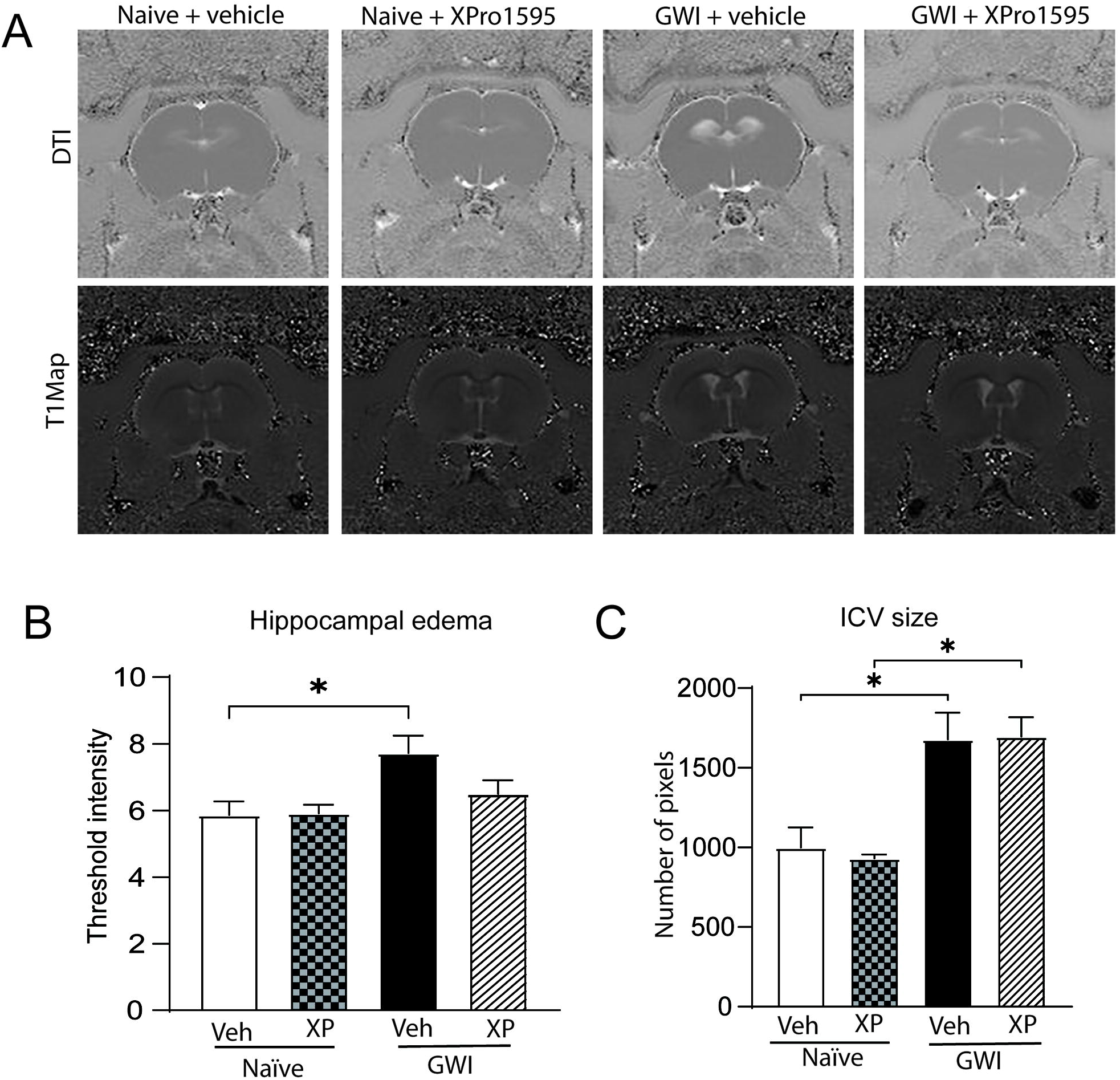
Magnetic Resonance Imaging T1-weighted (**A**) and Diffusion tensor imaging (DTI) was performed on the rat brains. Hippocampal level of intensity DTI slices (∼ bregma -3.14 mm) was semi-quantitated (**B**). The GWI+Veh group had a significant increase in intensity compared to the control group (Naïve+Veh), which was absent in the XPro1595-treated GWI group (GWI+XPro). The lateral ventricle size was quantitated in T1-MAP slices (∼ bregma -0.80 mm) (**C**). The GWI+Veh group had a significant increase in lateral ventricle size, compare to the Naïve+Veh group, that was not improved following XPro1595 treatment. *n* = 6 animals per group; * p = 0.05.

### GWI Increases Ventricle Size, which is not Attenuated by Neutralizing solTNF

Increased ventricular size is observed in Veterans with GWI, and this may be caused by inflammation and edema, therefore we wanted to determine whether DFP exposure increases ventricular size, and any impact of neutralizing solTNF. Using the T1 MRI scans we quantitated lateral ventricular size and observed that vehicle-treated GWI rats (GWI+Veh) had significantly larger ventricles than the control rats (Naïve+Veh) (Figure 5C). Two weeks of XPro1595 treatment had no effect on reducing the size of the lateral ventricle (GWI+XPro).

### Hippocampal Dendritic Spine Density and Morphology is Unaltered Chronically Following DFP Exposure

To begin determining the effects of altered neuroinflammation and edema on hippocampal dendritic plasticity, hippocampal CA1 dendritic spine density and morphology was quantitated. Spine density was quantitated on 200 µm lengths of both apical and basal dendrites (Figure 6A). Analysis revealed that DFP exposure does not increase spine density chronically (GWI+Veh) (Figure 6B), which remained unaffected following XPro1595 treatment (GWI+XPro). Next, we quantitated differences in spine morphology and did not observe any differences between groups (Figure 6C-H). To ensure a lack of effect across groups was not due to the location along the dendrite where dendritic density was determined, we quantitated the distance between the cell body and the start of the 200 µm length of dendrite that was used for analysis, and found no significant difference across groups (Figure 6H).

**Figure 6:**
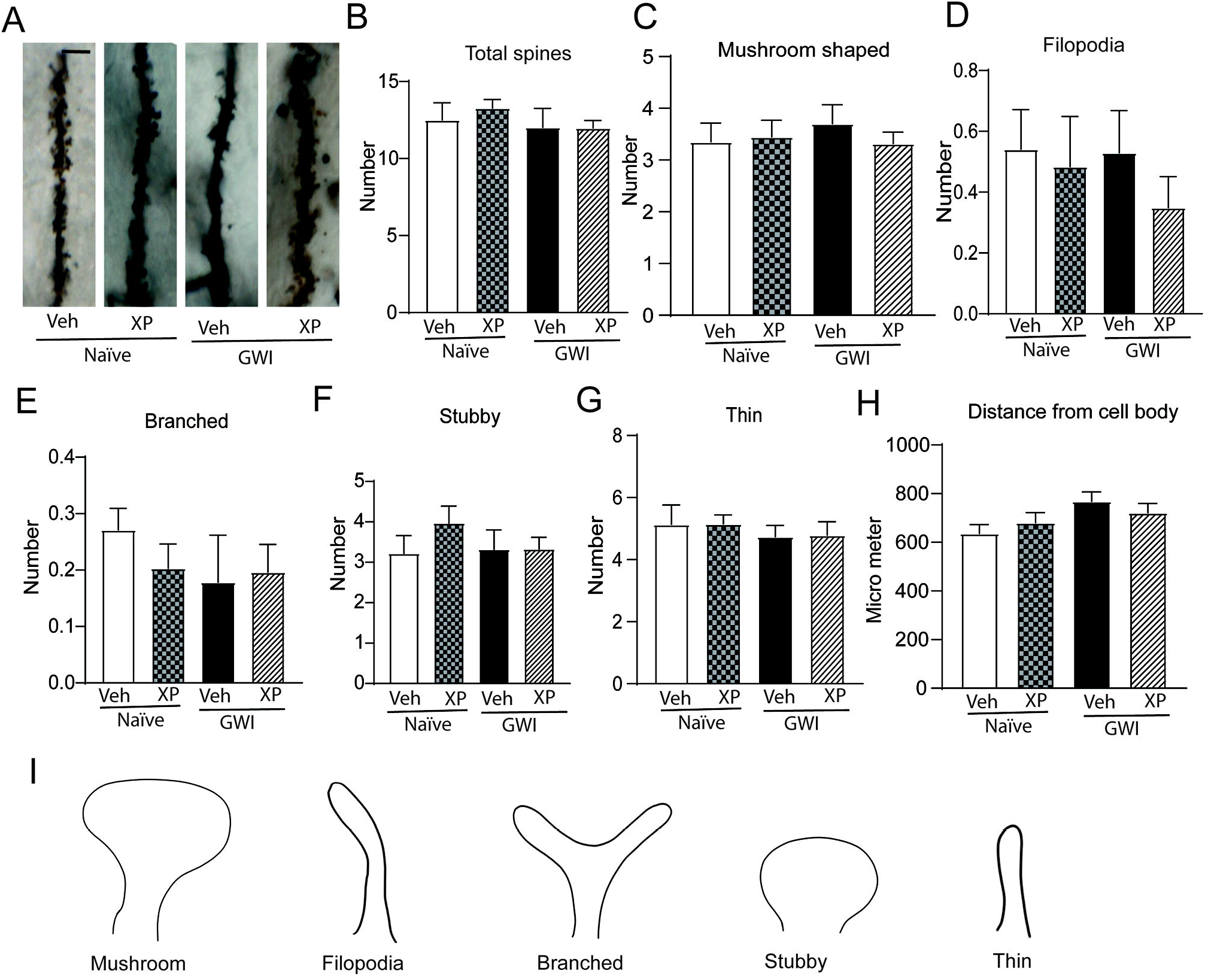
The effect of DFP exposure and subsequent XPro1595 treatment on hippocampal CA1 dendritic spine density (on a 200 µm segment of dendrite) and morphology was quantitated. No significant differences were observed between groups on any parameter measured. Representative Golgi stained neurons are shown in (**A**) for each experimental condition. Graphs show the total number of spines quantitated (**B**), mushroom-shaped spines (**C**), filopodia shaped spines (**D**), branched spines (**E**), stubby spines (**F**), and thin spines (**G**) analyzed, as well as distance of 200 μm dendritic segment from cell body (**H**). Examples of each spine category displayed in (**I**). *n* = 8 animals per group. Scale bar in A = 1 μm.

### GWI Impairs Post-Synaptic Protein Expression (Without Affecting Pre-Synaptic or Astrocytic Synaptic Protein Levels), which is Attenuated after Neutralizing solTNF

Neuroinflammation, specifically the inflammatory cytokine TNF, is a known regulator of synaptic plasticity (60). Therefore, to begin determining the role of hippocampal solTNF/TNFR1 activity in dendritic synaptic plasticity following chronic exposure to DFP, we quantitated major proteins associated with the tripartite synapse (pre-synaptic, post-synaptic and astrocytic glial component). There were no changes observed to pre-synaptic synaptophysin or astrocytic EAAT2 levels in the GWI rats (Figure 7 A&C). Conversely, there was a significant reduction in post-synaptic PSD-95 expression in the GWI+Veh group that was not present following XPro1595 treatment (GWI+XPro) (Figure 7B). Despites finding no alterations to dendritic spine density or morphology, a reduction in PSD-95 suggests there could be impaired trafficking of neurotransmitter receptors to and from the post-synaptic membrane.

**Figure 7:**
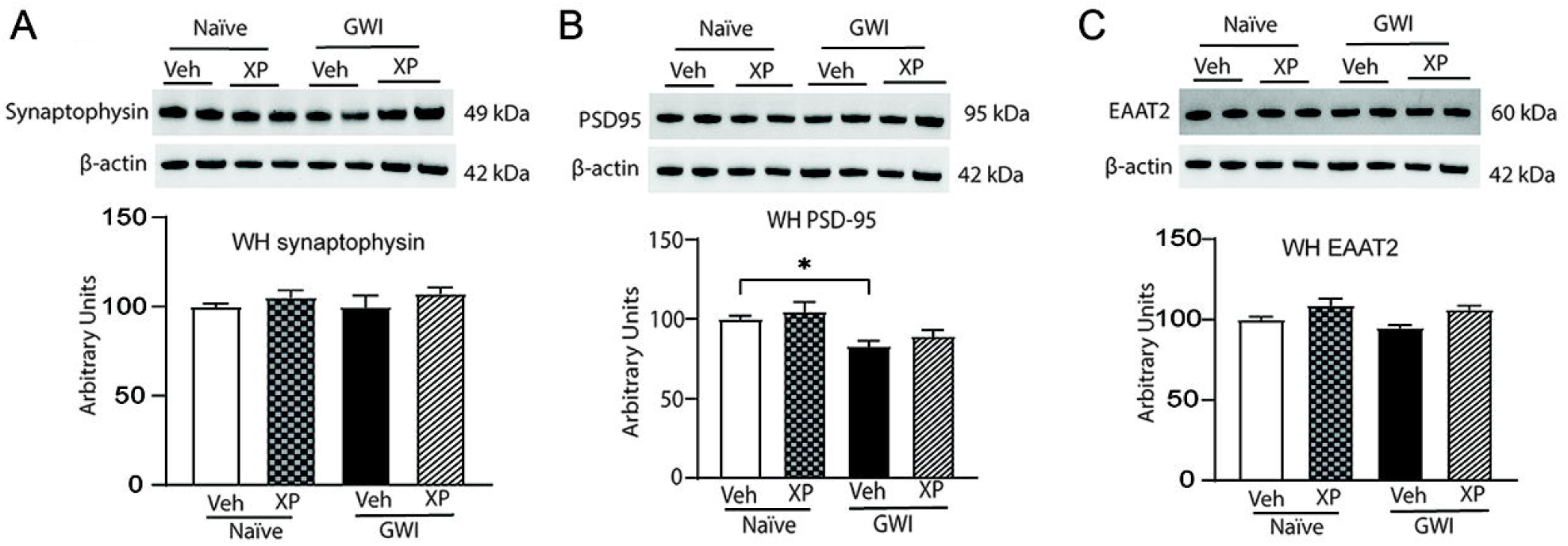
The hippocampus was dissected and processed for Western blotting to analyze pre-synaptic (synaptophysin), post-synaptic (PSD-95) and astrocytic (EAAT2) proteins involved in the tripartitate synapse. Quantification revealed no change in pre-synaptic (**B**) or astrocytic (**C**) protein expression between groups, however significant increases in PSD-95 expression in the GWI+Veh group (**A**) was observed, compared to the control group (Naïve+Veh), and which was absent following XPro1595 treatment (GWI+XPro). n=6 animals per group; * p = 0.05.

### GWI Impairs AMPAR1 Phosphorylation that is Not Improved By solTNF Neutralization

To determine alterations in neurotransmitter receptor activity, we first quantitated AMPAR1 and NMDAR1 receptor expression levels, and confirmed no changes in the total levels of hippocampal AMPAR1 and NMDAR1 receptors across the groups (Figure 8A&D). Conversely, because PSD-95 can regulate glutamate receptor phosphorylation that alters its ability to translocate to the post-synaptic membrane (61), we quantitated AMPAR1 and NMDAR1 receptor phosphorylation levels. Interestingly, we observed a significant reduction in p-AMPAR1 (Ser845) (Figure 8C) in the GWI+Veh group (tendency for reduced p-AMPAR1 (Ser831) (Figure 8B) and p-NMDAR1 (Ser896) expression) (Figure 8E), in addition to a significant reduction in all 3 phospho-antibodies in the GWI+XPro group (p-AMPAR1 (Ser845 and Ser831) and p-NMDAR1 (Ser896)).

**Figure 8:**
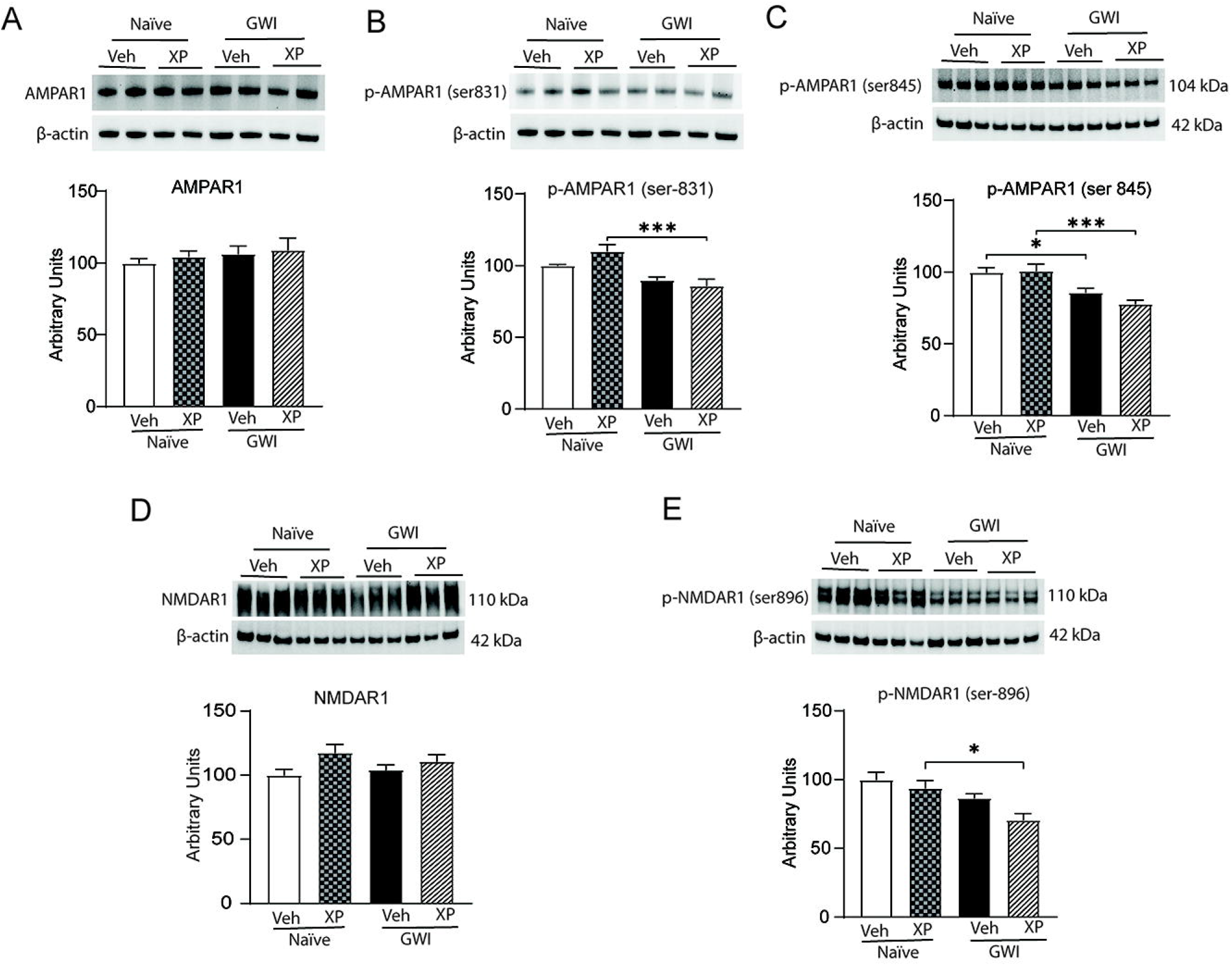
The hippocampus was dissected and processed for Western blotting to analyze post-synaptic neurotransmitter receptor expression and phosphorylation. Quantification revealed no changes in the total levels of hippocampal AMPA (**A**) and NMDA (**D**) receptors across the groups. AMPA receptor phosphorylation (**B&C**) and NMDA receptor phosphorylation (**E**) levels were also quantitated. A significant reduction was observed in p-AMPA receptor (Ser845) expression, along with a tendency for reduced p-NMDA receptor expression in the GWI+Veh group, which was not improved with XPro1595 treatment. n = 6 animals per group; * p = 0.05, *** p = 0.001.

### GWI Impairs Phosphorylated AMPAR Trafficking into the Post-Synaptic Membrane, which is Prevented by Neutralizing solTNF

Despite a lack of differences in the total level of neurotransmitter receptor phosphorylation across the two GWI groups (GWI+Veh and GWI+XPro), the reduced expression of PSD-95 in the GWI+Veh group suggests impaired trafficking of AMPAR1 and NMDAR1 receptors to the post-synaptic membrane. Therefore, we isolated the hippocampal synaptoneurosome to assess neurotransmitter receptor levels at the synaptic membrane. To first confirm the purity of our synaptoneurosome preparations, we compared PSD-95, synaptophysin, EAAT2 and Histone H3 expression in untreated rat whole hippocampus and synaptoneurosome preparations. We observed that the synaptoneurosome preparations were enriched for post-synaptic proteins (PSD-95) (Supp Figure S1A&B) without any nuclear (Histone H3) contamination Supp Figure S1A&E). While there appears to be an equal level of synaptophysin expression and a reduction in EAAT2 expression in the synaptoneurosome preparation compared to the whole hippocampus tissue (Supp Figure A), once beta-actin expression is accounted for in the synaptoneurosome samples the preparations also become enriched for synaptophysin and EAAT2 (Supp Figure C&D).

Using this synaptoneurosome preparation, a significant reduction was observed in the expression of PSD-95 in the GWI+Veh group (Figure 9A), compared to the control group, which was not improved by XPro1595 treatment. We next assessed total and phosphorylated AMPAR1 and NMDAR1 receptor levels at the synaptic membrane. Similar to whole hippocampal tissue, there were no changes in total AMPA receptor levels in the synaptic membrane (Figure 9B), however analysis of the ratio of AMPA receptor level/PSD-95 ratio reveals the GWI+Veh group had significantly increased AMPAR1 receptor levels once PSD-95 levels are normalized (Figure 9C), although this was not observed in the GWI+XPro group, suggesting that solTNF neutralization restores AMPAR1 receptor trafficking into the synaptic density. Similarly, we also observed a significant reduction in p-AMPAR1 (Ser 845) (Figure 9D) expression in the GWI+Veh group that was not present after neutralizing solTNF. No changes were observed in total or phosphorylated NMDAR1 receptor levels (Figure 9 E&F).

**Figure 9:**
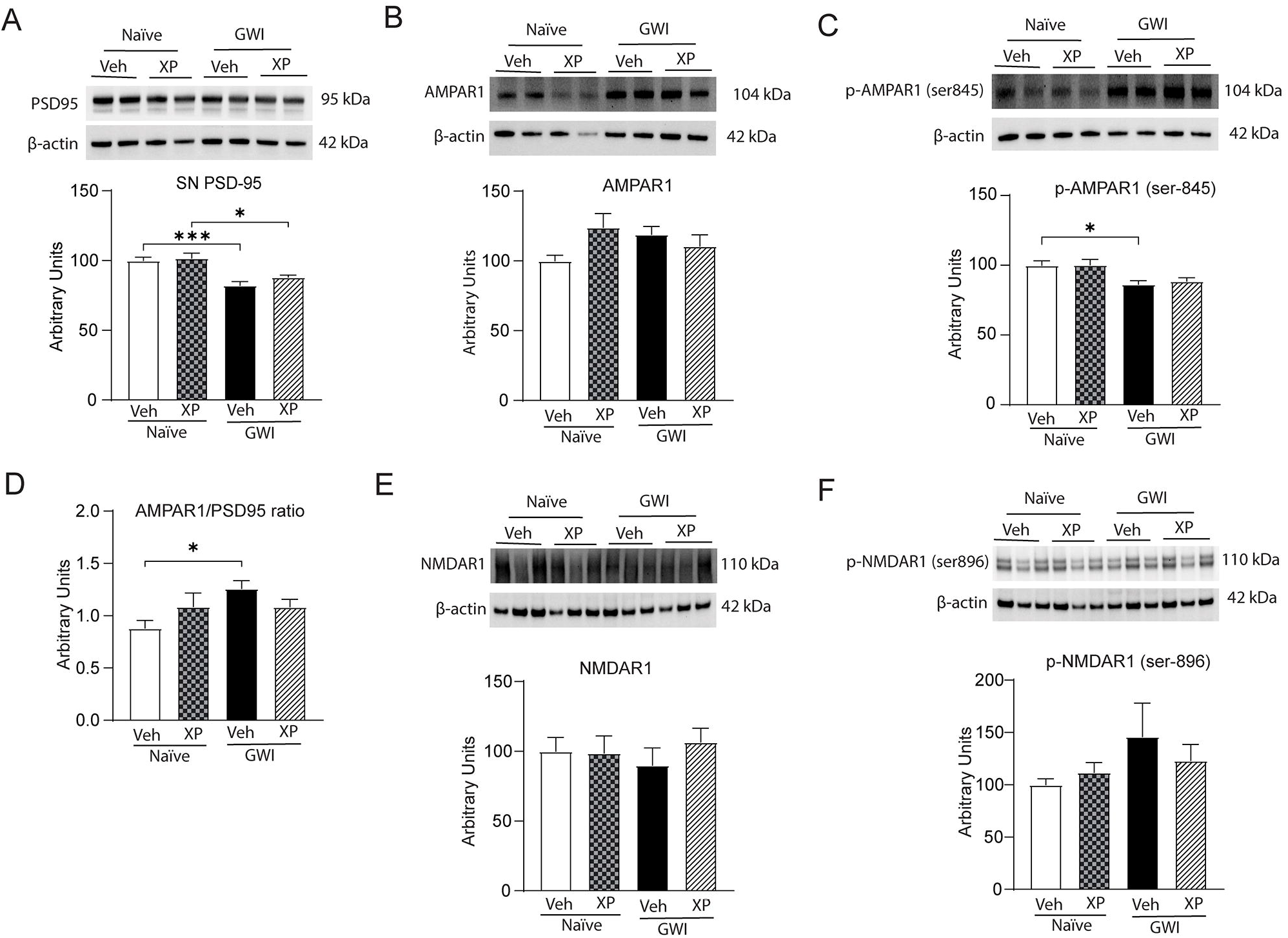
The hippocampus was dissected and the synaptoneurosome isolated prior to being processed for Western blotting to analyze post-synaptic protein expression. Quantification revealed a significant reduction in PSD-95 expression in the GWI+Veh group (**A**), which was not improved by XPro1595 treatment. Similar to whole hippocampal tissue, there were no changes in total AMPA receptor levels in the synaptic membrane (**B**), however analysis of the ratio of AMPA receptor level/PSD-95 ratio reveals the GWI+Veh group had significantly increased AMPA receptor levels once PSD-95 levels are normalized (**C**), which was not observed in the GWI+XP group. A significant reduction (**D**) in p-AMPAR (Ser 845) expression in the GWI+Veh group that was not present after neutralizing solTNF. No changes were observed in total or phosphorylated NMDA receptor levels (**E&F**). n = 6 animals per group; * p = 0.05, *** p = 0.001.

### Xpro1595 Attenuates DFP-Induced Chronic Cognitive Impairment by Neutralizing solTNF in GWI Rats

TNFR1-induced impairments in synaptic plasticity within the hippocampus are known to promote cognitive deficits (43). Therefore, we wanted to assess the extent of cognitive abilities in the rats using the NOR test, BM and MWM. In both the BM and MWM there were no significant differences in the ability of the rats to learn the tasks (Figure 10 A&D), suggesting that the GWI model does not impact the ability of these animals to learn. However, in all three of the cognitive tests the GWI+Veh rats were significantly less able to remember the task at a later time point (BM and MWM = 4 days; NOR = 1 hour) (Figure 10 B,C,E-H). In the BM the GWI+Veh rats were less able to remember where the target hole was located (Figure 10 B&C), in the MWM they spent less time in the platform quadrant (Figure 10E), and in the NOR test they spent significantly less time with the novel object (Figure 10 F-H). Conversely, the data from all three cognitive tests show that the GWI+XPro group had become significantly better than the GWI+Veh group, and were no longer significantly worse than the Naïve+XPro group.

**Figure 10:**
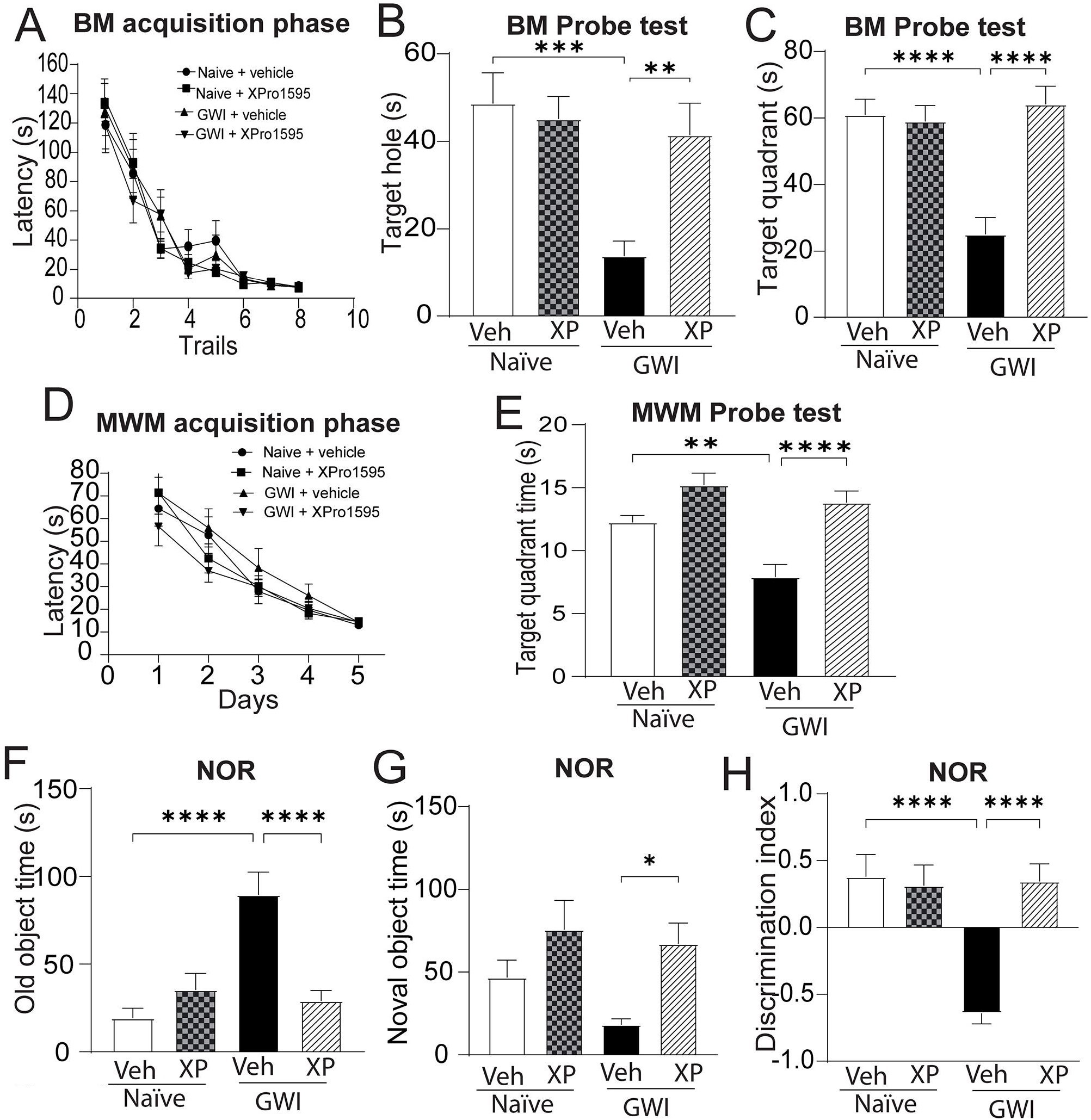
Cognitive abilities of the rats were assessed using the Novel Object Recognition (NOR) test, Barnes Maze (BM) and Morris Water Maze (MWM). No significant differences were observed in the abilities of the rats to learn on the MWM and BM (**A&D**). However, in each of the cognitive tests the GWI+Veh rats were significantly less able to remember the task at a later time point (MWM and BM = 4 days; NOR = 1 hour). In the NOR test the GWI+Veh rats spent significantly less time with the novel object (**F&H**) and more time with the old object (**G**), in the BM they were less able to remember where the target hole was located (**B&C**), and in the MWM they spent less time in the platform quadrant (**E**). Conversely, the data from all 3 tests show that rats from the GWI+XPro group had become significantly better than the GWI+Veh group, and were no longer significantly worse than rats from the Naïve+XPro group. For each cognitive test, Naïve+Veh (n=12), Naïve+XPro (n=11), GWI+Veh (n=12), GWI+XPro (n=13) animals per group; * p = 0.05, ** p = 0.01, *** p = 0.001, **** p = 0.0001.

### Xpro1595 Attenuates Anxiety-Like Behavior thru Neutralizing solTNF in GWI Rats

TNF has been shown to regulate hippocampal synaptic plasticity to modulate anxiety related behavior (62), therefore we wanted to assess the anxiety-like behavior of the rats using the MB, OF and EPM tests. In the MB, OF and EPM tests the GWI+Veh group spent significantly more time burying marbles (Figure 11A), less time in the OF center zone (Figure 11B) and less time and entries in the open arms (Figure 11C-E), suggesting DFP exposure promotes anxiety. Conversely, treating the GWI rats with XPro1595 (GWI+XPro group) diminished their anxiety-like behavior, so that they were no longer significantly different to the control groups.

**Figure 11:**
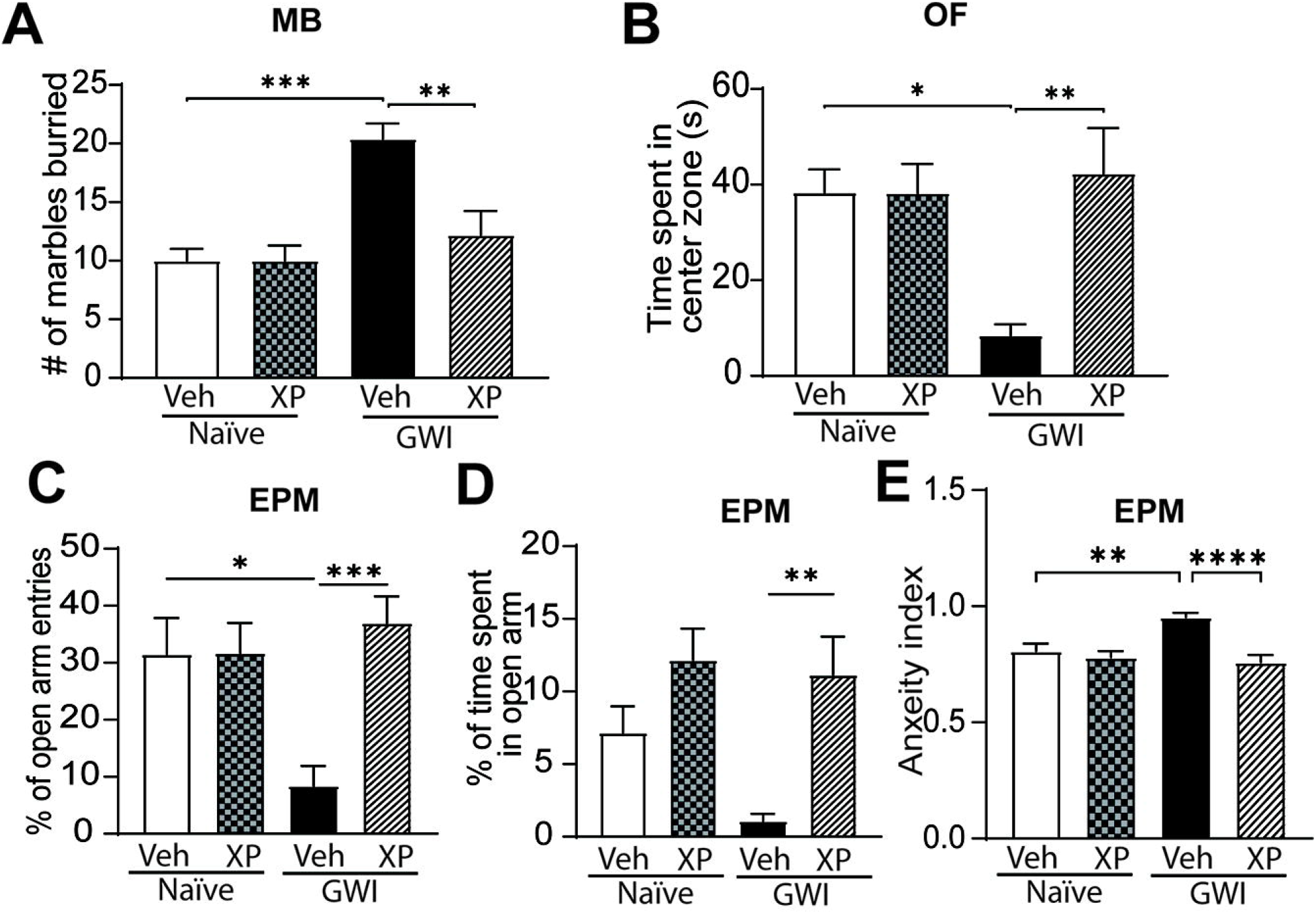
Anxiety behaviors were assessed using the Marble Burying (MB) test, Open Field (OF) and Elevated Plus Maze (EPM). In the MB test the rats in the GWI+Veh group spent significantly more time burying marbles (**A**), compared to the control groups, as reflected by more marbles having been buried in the same amount of time. Similarly, GWI+Veh rats spent significantly less time in the center zone of the OF arena (**B**) and in the EPM open arms (**C-E**), collectively suggesting that these animals have increased anxiety. Conversely, treating the GWI rats with XPro1595 (GWI+XPro group) diminished their anxiety-like behavior, so that they were no longer significantly different to the control groups. For the MB test: Naïve+Veh (n=5), Naïve+XPro (n=5), GWI+Veh (n=5), GWI+XPro (n=5) animals per group. For the OF and EPM: Naïve+Veh (n=17), Naïve+XPro (n=16), GWI+Veh (n=17), GWI+XPro (n=18) animals per group. * p = 0.05, ** p = 0.01, *** p = 0.001, **** p = 0.0001.

### GWI Promotes Anhedonia and Neuropathic Pain that is Attenuated After Neutralizing solTNF by XPro1595

TNFR1-induced impairments in synaptic plasticity within the hippocampus are known to cause depressive-like behavior and hindpaw mechanical hypersensitivity (63)(31)(32). Therefore, we wanted to determine whether the altered inflammatory response and synaptic plasticity in the GWI animals promotes anhedonia and neuropathic pain. We observed that the GWI+Veh group drank significantly less sucrose water in the SPT, (Figure 12A), and had more hindpaw hypersensitivity (Figure 12B), compared to control rats, suggesting that the GWI rats display anhedonia and neuropathic pain. Conversely, once the rats were treated with XPro1595 (GWI+XPro) these behaviors were no longer significantly different to the control rats.

**Figure 12:**
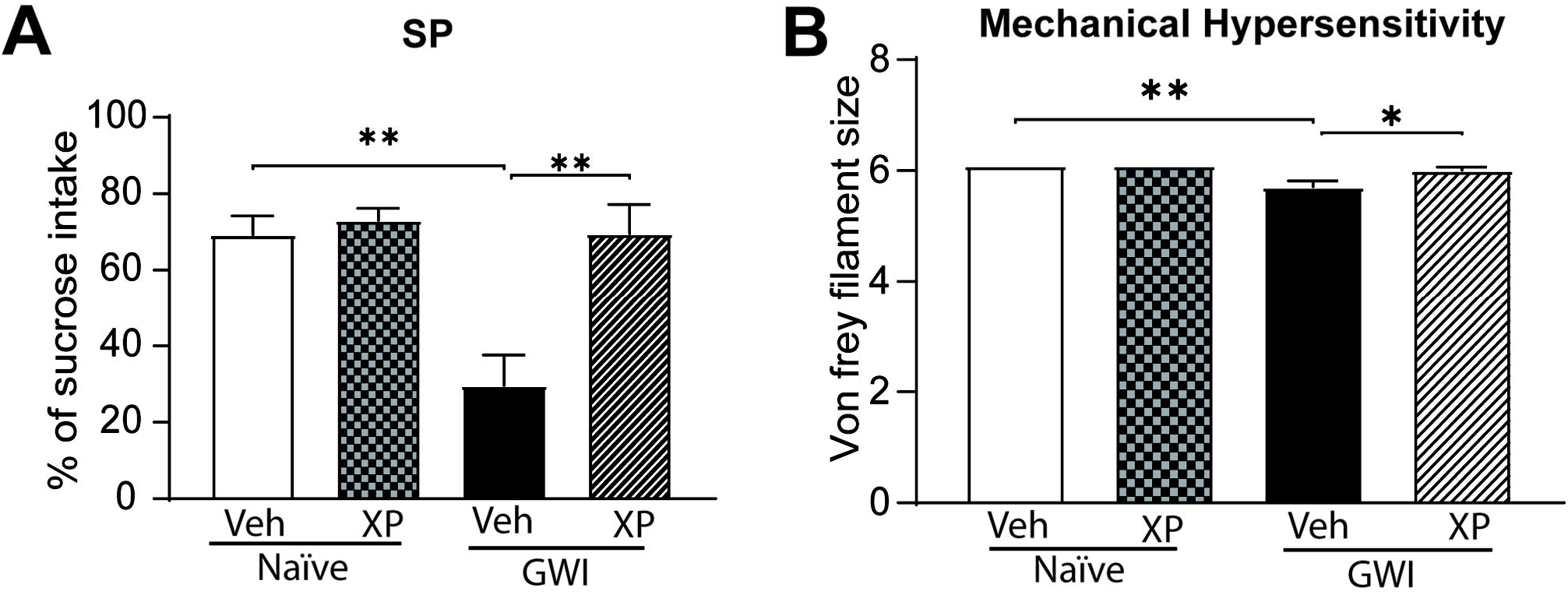
Depressive-like behaviors and neuropathic pain were assessed using the Sucrose Preference Test (SP) and Hindpaw mechanical hypersensitivity. In the SP test rats in the GWI+Veh group drank significantly less sucrose water (**A**), and had more hindpaw hypersensitivity (**B**), compared to control rats (Naive+Veh), suggesting that the GWI rats display anhedonia and neuropathic pain. Conversely, once the rats were treated with XPro1595 these behaviors were no longer significantly different to the control rats. For both tests (*n* = 5) animals per group. * p = 0.05, ** p = 0.01.

## DISCUSSION

### GWI Impairs Neurotransmitter Receptor Trafficking Mechanisms

Veterans with GWI experience a cluster of medically unexplained chronic symptoms including mood and memory impairment (64), and is confirmed in many animal models with impairments in memory observed, as well as anxiety, depressive-like behavior and neuropathic pain (65), (66) (10) (67) (68) (69). Data from the current study confirm these findings, but also begins to unravel intracellular mechanisms of action, whereby DFP exposure impairs the cellular mechanisms responsible for trafficking AMPA receptors to the synaptic membrane. Under normal physiological conditions cholinergic activity promotes glutamatergic activity (52) (70), and therefore impairments to AMPA receptor trafficking mechanisms (altered PSD-95 expression levels) are not surprising following DFP exposure. Reductions in PSD-95 expression can then modulate AMPA receptor phosphorylation (61), which can disrupt neural circuitry function to impair neurological outcomes. Given that targeted mutations of the noncatalytic region of acetylcholinesterase can reduce PSD-95 expression (71), it is plausible to suggest that the DFP-induced irreversible acetylcholinesterase inhibition could be directly responsible for the observed reduction in PSD-95 expression in the current study, although further investigation is required.

### solTNF Inhibition Reduces Inflammation and Restores AMPA Receptor Trafficking Mechanisms

Recent GWI animal models also suggest neuroinflammation may underscore the observed neurological dysfunctions, including dysregulated reactivity of astrocytes and microglia (10) (6) (72) (73) (74) (75) (76), that contribute to an elevated cytokine expression (12). Studies in patients with GWI have also found autoantibodies against astrocytic, oligodendrocyte, and neuronal proteins (77), and elevated serum cytokine and cytokine receptor levels (78) (79), confirming a dysregulated inflammatory response. The elevated cytokine expression and glial reactivity observed chronically following DFP exposure in the current study adds further evidence of a neuroinflammatory component to the GWI condition. However, the connection between an elevated inflammatory response and impaired neurological function remained under studied.

Inflammation is a key regulator of synaptic plasticity (80), specifically TNFR1, which plays a major role in maintaining hippocampal homeostatic synaptic plasticity (regulation of AMPA receptor surface expression) under physiological conditions, which subsequently regulates mEPSC (excitatory postsynaptic potential) currents and eventually LTP (81), (30). Under pathological conditions injury (or simply TNF administration) promotes phosphorylation and translocation of AMPA receptors to the synaptic membrane within the first hour (82) (83), that may predispose the neuron to glutamate-mediated excitotoxicity leading to cell death. However, this is followed by a second phase that includes reductions in dendritic spine density and length (84) (31), and reduced post-synaptic protein expression (PSD-95) (85) (86), concomitant with a reduction in surface expression and synaptic localization of AMPA receptors (87) (88) to suppresses LTP (89) and thus modulate neural connectivity and animal behavior. Importantly, there is a strong connection between TNF/TNFR1-mediated synaptic plasticity, and the development of impaired neurological outcomes (63) (90) (91) (92). In the current study XPro1595 neutralization of solTNF overcame DFP-induced impairments to neurotransmitter trafficking mechanisms sufficiently enough to improve restore cognition, in addition to reducing anxiety, depressive-like behaviors and neuropathic pain. Intriguingly however, the total level of AMPAR1 observed within the synaptoneurosome fraction was unaltered between groups, and therefore there are questions remaining to be answered. Since TNF can promote insertions of GluA2-lacking AMPARs into the plasma membrane (93), then future studies should consider evaluating the impact of the presence or absence of GluA2 in this system.

An intriguing question that arises from these studies is how using XPro1595 to neutralize solTNF may be reversing DFP-induced impairments, and thus is there any clinical relevance? While rodent studies show increased hippocampal and serum TNF expression acutely and 6 weeks following GWI induction (11) (94), studies on Veterans in the chronic environment with diagnosed GWI show baseline serum TNF levels (95) (96), and our studies also show no difference in TNFR1 protein expression chronically following DFP exposure. However, even small variances in TNF expression that may be difficult to detect have been shown to drive pathological changes (29). One possible overlap between DFP, TNFR1 and neurotransmitter receptor trafficking may be the JAK/STAT pathway. The JAK/STAT pathway has been shown to be involved in synaptic plasticity by way of NMDA activation (97), which subsequently interacts with downstream effectors of AMPA receptor trafficking, including PSD-95 (98). Importantly, both DFP and TNFR1 can activate JAK/STAT signaling (11) (99) making it plausible to suggest that inhibition of TNFR1 using XPro1595 could be modifying JAK/STAT regulation of neurotransmitter trafficking at the post-synaptic density, however further investigation is certainly required.

## CONCLUSION

Our study demonstrates that DFP exposure produces inflammation, edema, and abnormal synaptic plasticity associated with chronic neurological function impairment, like those commonly reported in GWI Veterans. Our study provides novel evidence that persistent cognitive dysfunction and chronic neuroinflammation in a model of GWI are linked with solTNF and its receptor TNFR1-mediated neuroinflammation in the hippocampus. Our results suggest that selectively inhibiting solTNF using XPro1595 reduces neuroinflammation, and improves neuroplasticity, and overall behavioral function when administered in the chronic setting of a rat model of GWI. Our findings support the use of XPro1595 in Veterans with GWI, and provide evidence that selective anti-inflammatory treatment targeting abnormal solTNF-mediated neuroinflammatory responses may be a good therapeutic strategy for GWI.

## Abbreviations

AMPA: α-amino-3-hydroxy-5-methyl-4-isoxazolepropionic acid
AMPAR1: α-amino-3-hydroxy-5-methyl-4-isoxazolepropionic acid receptor 1
p-AMPAR1: phospho-α-amino-3-hydroxy-5-methyl-4-isoxazolepropionic acid receptor 1
BBB: Blood-brain barrier
ANOVA: Analysis of variance
BM: Barnes Maze
DFP: Diisopropyl fluorophosphate
DG: Dentate gyrus
DTI: Diffusion tensor imaging
EAAT2: Excitatory amino acid transporter 2
EPM: Elevated plus maze
GFAP: Glial fibrillary acidic protein
GWI: Gulf war illness
Iba1: Ionized calcium binding adaptor molecule 1
LTD: Long-term depression
LTP: Long-term potentiation
MP: Marble burying
MRI: Magnetic resonance imaging
MWM: Morris water maze
NMDA: N-methyl-D-aspartate
NMDAR1: N-methyl-D-aspartate receptor 1
p-MNDAR1: phospho- N-methyl-D-aspartate receptor 1
NOR: Novel object recognition
PSD-95: Postsynaptic density protein 95
solTNF: Soluble tumor necrosis factor
SPT: Sucrose preference test
TNF: Tumor necrosis factor
TNFR1: Tumor necrosis factor receptor 1

## ACKNOWLEDGMENT

We acknowledge the funding and support of the Department of Defense GW190083 (KJD), the Virginia Department for Aging and Rehabilitative Services FP00001476 (KJD). Microscopy was performed in the VCU Microscopy Core (supported in part by funding from the NIH-NCI Cancer Center Support Grant P30 CA016059). We thank INmuneBio Inc. for their generous donation of XPro1595.

## AUTHOR CONTRIBUTIONS

KJD and LSD conceptualized the research; LSD developed the GWI model; KJD and UJ designed the experiments; UJ, KL, NNL, MD, AB, and EH performed the experiments and collected the data; UJ analyzed the data; UJ and KJD interpreted the data and wrote the manuscript.

## FIGURE LEGEND

**Supplementary Table 1: List of antibodies used in the study**.

**Supplementary Figure 1:** The hippocampus from naïve animals was dissected and processed for synaptoneurosome isolation prior to being processed for Western blotting to confirm purity of the isolation technique. Samples were analyzed for pre-synaptic (synaptophysin), post-synaptic (PSD-95), astrocytic (EAAT2) tripartite synapses, as well as for any nuclear contamination (Histone-H3). Synaptoneurosome samples were compared to whole hippocampal samples (**A**). Synaptoneurosome samples were enriched for PSD-95 (**B**), without any nuclear Histone-H3 (**E**) contamination. While there appears to be an equal level of synaptophysin expression (**C**) and a reduction in EAAT2 expression (**D**) in the synaptoneurosome preparation compared to whole hippocampal tissue, once beta-actin is normalized the samples become enriched for both synaptophysin and EAAT2. n = 2 animals per group.

