## Supplemental Table S1 for "SELECTIVE INHIBITION OF SOLUBLE TNF ATTENUATES HIPPOCAMPAL NEUROINFLAMMATION AND PSD-95 EXPRESSION TO IMPROVE NEUROLOGICAL FUNCTIONS IN A RAT MODEL OF GULF WAR ILLNESS"

| Primary antibodies | | | | | | | | | | |
| --- | --- | --- | --- | --- | --- | --- | --- | --- | --- | --- |
| Anti- | | | Abbreviation | Host | | Company | Catalog number | Application | | Dilution ^a^ |
| Glial fibrillary acidic protein | | | GFAP | Rabbit | | Dako | Z0334 | IHC | | 1:200 |
| Ionizing calcium-binding adaptor molecule 1 | | | IBA-1 | Rabbit | | Wako | 01919741 | IHC | | 1:200 |
| Excitatory Amino Acid Transporter-2 | | | EAAT-2 | HRP-linked | | Santa Cruz | Sc-365634HRP | WB | | 1:1000 |
| Postsynaptic density protein 95 | | | PSD-95 | Mouse | | NeuroMab | - | WB | | 1:200 |
| Tota l- α-amino-3-hydroxy-5-methyl-4-isoxazole propionic acid receptor 1 | | | t-AMPAR1 | Rabbit | | Millipore | AB1504 | WB | | 1:1000 |
| Phospho - α-amino-3-hydroxy-5-methyl-4-isoxazolepropionic acid receptor 1 (ser831) | | | p-AMPAR1 (ser831) | Rabbit | | Millipore | 04823 | WB | | 1:1000 |
| Phospho - α-amino-3-hydroxy-5-methyl-4-isoxazolepropionic acid receptor 1 (ser845) | | | p-AMPAR1 (ser845) | Rabbit | | R&D System | PPS008 | WB | | 1:1000 |
| Total - N-methyl-D-aspartate receptor 1 | | | t-NMDAR1 | Rabbit | | Millipore | MAB1586 | WB | | 1:1000 |
| Phospho - N-methyl-D-aspartate receptor1 (ser896) | | | p-NMDAR1 (ser896) | Rabbit | | Invitrogen | PA537589 | WB | | 1:1000 |
| Synaptophysin | | | SYP | Mouse | | Millipore | MAB368 | WB | | 1:1000 |
| Tumor necrosis factor-α | | | TNF-α | Mouse | | Santa Cruz | sc-133192 | WB | | 1:200 |
| Tumor necrosis factor receptor 1 | | | TNFR1 | Rabbit | | Santa Cruz | sc-8436 | WB | | 1:1000 |
| Tumor necrosis factor receptor 2 | | | TNFR2 | Rabbit | | Santa Cruz | sc-8041 | WB | | 1:1000 |
| β-actin | | | β-actin | HRP-linked | | Santa Cruz | sc-47778HRP | WB | | 1:3000 |
| Secondary antibodies | | | | | | | | | | |
| Anti- | Host | Label | | | Catalog number | | Application | Dilution | Company | |
| Rabbit IgG | Goat | HRP-linked | | | 7074S | | WB | 1:3000 | Santa Cruz | |
| Mouse IgG | Horse | HRP-linked | | | 7076S | | WB | 1:3000 | Santa Cruz | |
| Rabbit IgG | Goat | Alexa 488 | | | A11034 | | IHC | 1:500 | Invitrogen | |

List of primary and secondary antibodies used in the study.

WB = Western blot; IHC = Immunohistochemistry

^a^ Dilutions were from original stocks supplied by the vendors.
