## Supplementary figures and images for "SELECTIVE INHIBITION OF SOLUBLE TNF ATTENUATES HIPPOCAMPAL NEUROINFLAMMATION AND PSD-95 EXPRESSION TO IMPROVE NEUROLOGICAL FUNCTIONS IN A RAT MODEL OF GULF WAR ILLNESS"

### Supplemental Figure S1

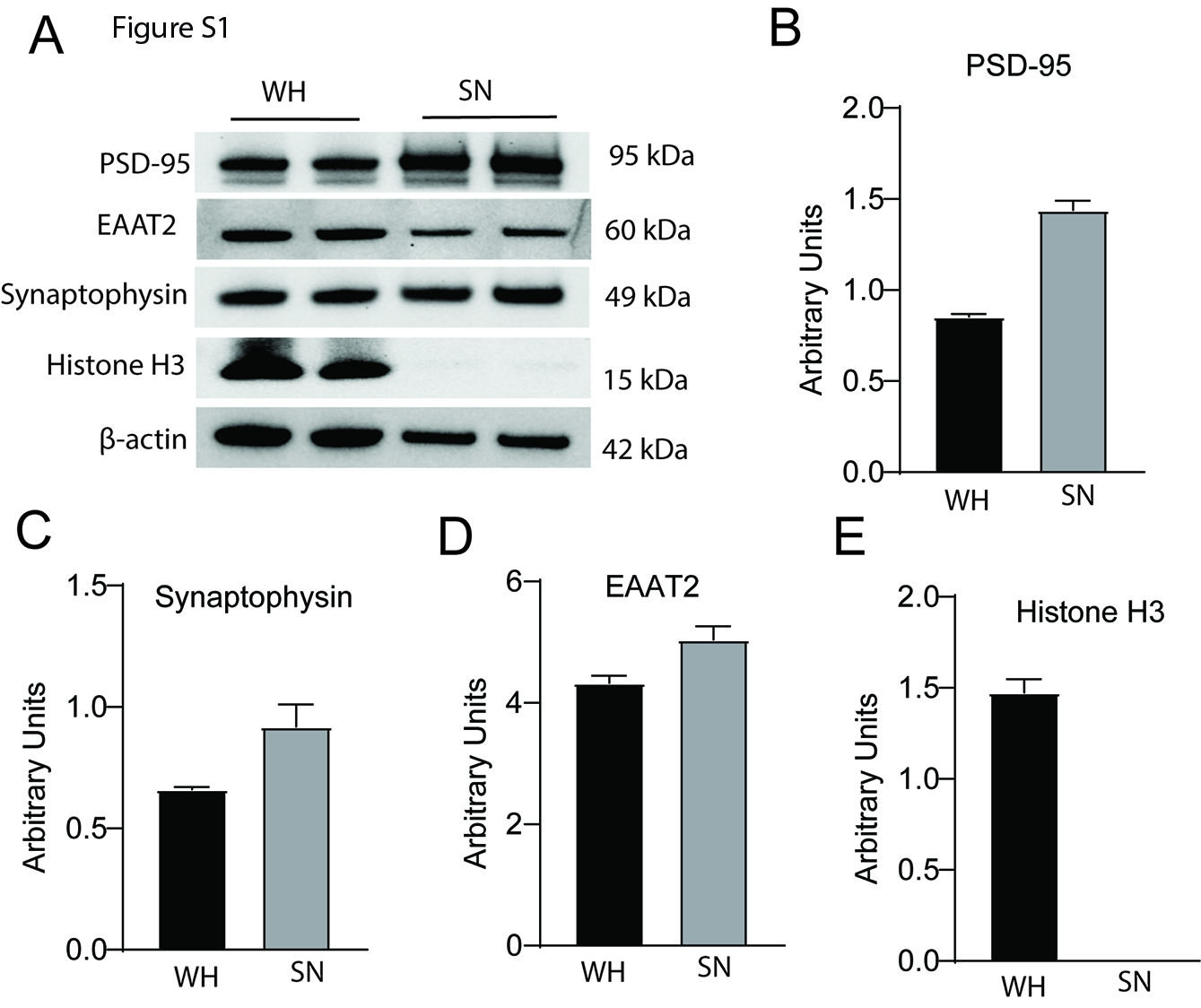
